# The structural basis for release factor activation during translation termination revealed by time-resolved cryogenic electron microscopy

**DOI:** 10.1101/470047

**Authors:** Ziao Fu, Gabriele Indrisiunaite, Sandip Kaledhonkar, Binita Shah, Ming Sun, Bo Chen, Robert A. Grassucci, Måns Ehrenberg, Joachim Frank

## Abstract

When the mRNA translating ribosome encounters a stop codon in its aminoacyl site (A site), it recruits a class-1 release factor (RF) to induce hydrolysis of the ester bond between peptide chain and peptidyl-site (P-site) tRNA. This process, called termination of translation, is under strong selection pressure for high speed and accuracy. Class-1 RFs (RF1, RF2 in bacteria, eRF1 in eukarya and aRF1 in archaea), have structural motifs that recognize stop codons in the decoding center (DC) and a universal GGQ motif for induction of ester bond hydrolysis in the peptidyl transfer center (PTC) 70 Å away from the DC. The finding that free RF2 is compact with only 20 Å between its codon reading and GGQ motifs came therefore as a surprise^1^. Cryo-electron microscopy (cryo-EM) then showed that ribosome-bound RF1 and RF2 have extended structures^2,3^, suggesting that bacterial RFs are compact when entering the ribosome and switch to the extended form in a stop signal-dependent manner^3^. FRET^4^, cryo-EM^5,6^ and X-ray crystallography^7^, along with a rapid kinetics study suggesting a pre-termination conformational change on the millisecond time-scale of ribosome-bound RF1 and RF2^8^, have lent indirect support to this proposal. However, direct experimental evidence for such a short-lived compact conformation on the native pathway to RF-dependent termination is missing due to its transient nature. Here we use time-resolved cryo-EM^9,10,11,12,13^ to visualize compact and extended forms of RF1 and RF2 at 3.5 and 4 Å resolution, respectively, in the codon-recognizing complex on the pathway to termination. About 25% of ribosomal complexes have RFs in the compact state at 24 ms reaction time after mixing RF and ribosomes, and within 60 ms virtually all ribosome-bound RFs are transformed to their extended forms.

## Main

Most intracellular functions are carried out by proteins, assembled as chains of peptide-bond linked amino acid (aa) residues on large ribonucleoprotein particles called ribosomes. The aa-sequences are specified by information stored as deoxyribonucleic acid (DNA) sequences in the genome and transcribed into sequences of messenger RNAs (mRNAs). The mRNAs are translated into aa-sequences with the help of transfer RNAs (tRNAs) reading any of their 61 aa-encoding ribonucleotide triplets (codons). In termination of translation, the complete protein is released from the ribosome by a class-1 release factor (RF) recognizing one of the universal stop codons (UAA, UAG, and UGA), signaling the end of the amino acid encoding open reading frame (ORF) of the mRNA. There are two RFs in bacteria, RF1 and RF2, one in eukarya, eRF1, and one in archaea, aRF1.RF1 and RF2 read UAA, UAG, and UAA, UGA, respectively, while the omnipotent eRF1 and aRF1 factors read all-codons. Stop codon-reading by RFs is aided by class-2 RFs, the GTPases RF3, eRF3 and aRF3 in bacteria, eukarya and, archaea, respectively. Each stop codon in the decoding center (DC) is recognized by a stop-codon recognition (SCR) motif in a class-1 RF, and all RFs have a peptidyl transfer center (PTC)-binding GGQ motif, named after its universal Gly-Gly-Glu triplet (GGQ), for coordinated ester bond hydrolysis in the P-site bound peptidyl-tRNA.

The crystal structures of free RF1 and RF2 have a distance between the SCR and GGQ motifs of about 20 Å^1,14^, much shorter than the 70 Å separating DC and PTC. This separating distance made the expected coordination between SCR and ester bond hydrolysis enigmatic. The crystal structure of free eRF1 has, in contrast, about 70 Å between its SCR and GGQ motifs, a distance close to the 80 Å between the DC and PTC of the 80S ribosome in eukarya^15^. Further cryo-EM work showed that ribosome-bound RF1 and RF2 have extended structures^2,3^, facilitating coordinated codon recognition in DC and ester bond hydrolysis in PTC. Subsequent high-resolution X-ray crystal^7,16,17,18,19,20,21,22^ and cryo-EM^5,6,23,24,25,26^ structures of RF-bound 70S ribosomes allowed the modeling of stop-codon recognition by RF1, RF2^27^, eRF1^28^ and GGQ-induction of ester bond hydrolysis^28^.

If the compact forms of free RFs in the crystal^1,14^ are physiologically relevant, it would mean that eubacterial RFs are in the compact form upon A-site entry (*pre-accommodation* state) and assume the extended form (*accommodation* state) in a stop-codon dependent manner. The relevance is indicated by a compact crystal structure of RF1 in a functional complex with its GGQ-modifying methyltransferase^29,30^, although SAXS data indicated free RF1 to be extended in bulk solution^31^. The existence of such a conformational switch would make high-resolution structures of these compact and transient RF-forms necessary for a correct description of the stop-codon recognition process hitherto based on post-termination ribosomal complexes^32,33^.

Indirect evidence for rapid conformational activation of RF1 and RF2 after A-site binding has been provided by quench-flow based kinetics^8^, and in a series of recent FRET experiments Joseph and collaborators showed free RF1 to be in a compact form^4^, compatible with the crystal forms of RF1^1^ and RF2^14^, but in an extended form when A-site bound to the stop-codon programmed ribosome^4^. Ribosome-bound class-1 RFs in the compact form has been observed together with alternative ribosome-rescue factor A (ArfA) in ribosomal rescue complexes, which lack any codon in the A site^5,6^. Very recently, Korostelev and collaborators (2018)^7^ used X-ray crystallography in conjunction with the peptidyl transfer-inhibiting antibiotic blasticidin S (Bl-S) to capture a mutated, hyper-accurate variant of RF1 in the stop codon-programmed termination complex. They found RF1 in a compact form, which they used to discuss stop-codon recognition in conjunction with large conformational changes of the RFs. It seems, however, that this Bl-S-halted ribosomal complex is in a post-recognition state (*i.e*., stop-codon recognition motif has the same conformation as in the post-accommodation state in DC) but before RF-accommodation in the A site, making its relevance for on-pathway stop-codon recognition unclear.

Here, in contrast, we use time-resolved cryo-EM^9,10,11,12,13^ for real-time monitoring of how RF1 and RF2 ensembles change from compact to extended RF conformation in the first 100 ms after RF-binding to the pre-termination ribosome. These compact RF-structures, originating from short-lived ribosomal complexes previously out of reach for structural analysis, are seen at near-atomic resolution (3.5 - 4 Å). The time-dependent ensemble changes agree qualitatively with accompanying (Fig. 1) and previous^8^ quench-flow studies. We discuss the role of the compact structures of RF1 and RF2 for fast and accurate stop-codon recognition in translation termination.

**Figure 1.**
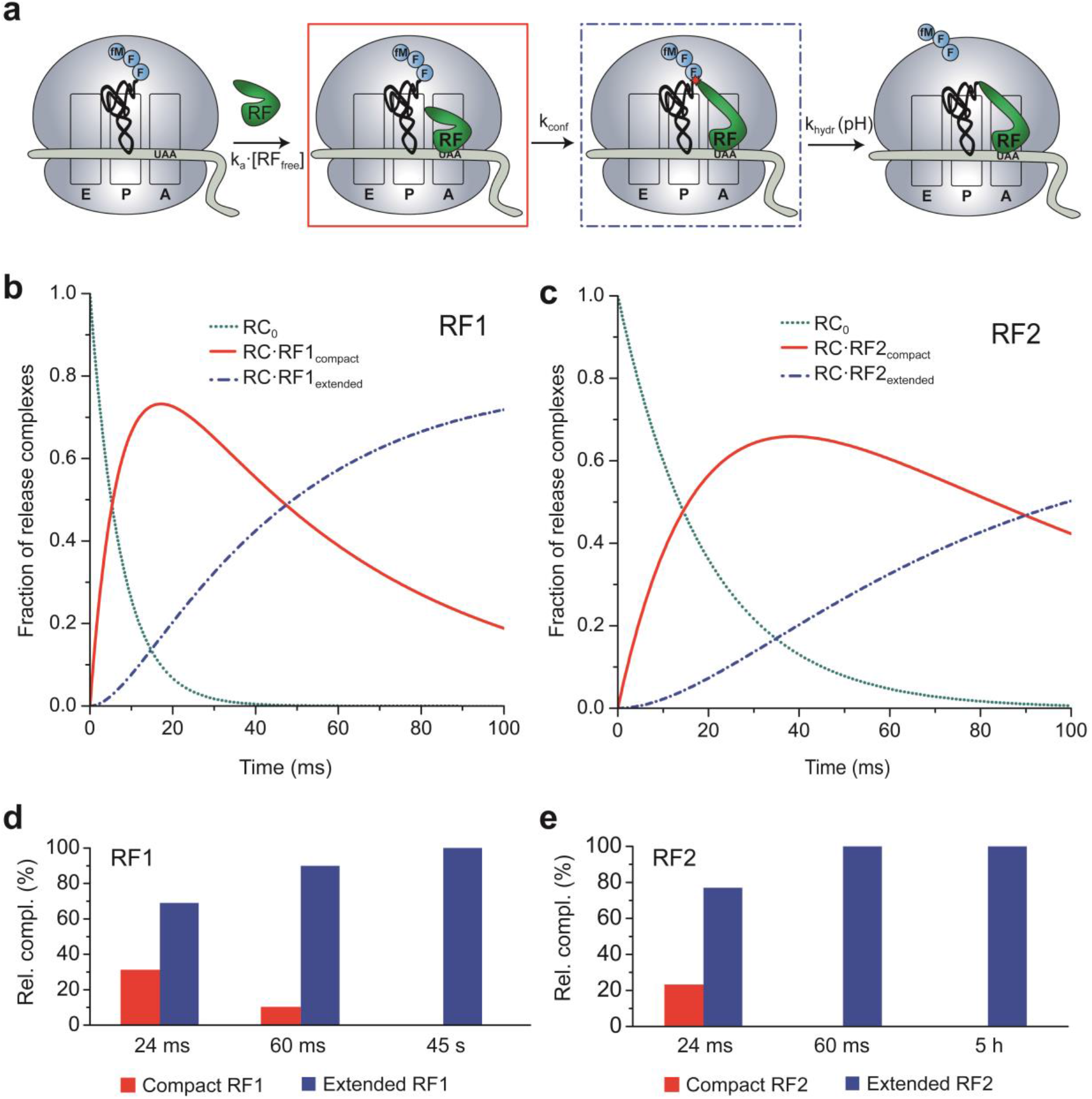
Time evolution of ribosome ensembles in termination of translation. **a.** Cartoon visualization of the pathway from free release complex to peptide release. Compact class 1 release factor (RF) binds to RF-free ribosomal release complex (RC_0_) and forms the RC·RFcompact complex with compounded rate constant k_a·_[RF_free_]. Stop codon recognition induces conformational change in the RF which brings the ribosome from the RC·RFcompact to the RC·RFextended complex with rate constant k_conf_. The ester bond between the peptide and the P-site tRNA is hydrolyzed with rate constant khydr. **b.** Predicted dynamics of peptide release with conformational change in RF1. We solved the ordinary differential equations associated with termination according to the scheme in Fig. 1a with association rate constant k_a_ = 45 µM^-1^s^-1^, [RF1_free_] = 3 µM, k_conf_ = 18 s^-1^ and k_hydr_ = 2 s^-1^ (Extended Data Fig. 2) and plotted the fractions of ribosomes in RC_0_, RC·RFcompact and RC·RFextended forms (y-axis) as functions of time (x-axis). **c.** Predicted dynamics of peptide release with conformational change in RF2. The fractions of ribosomes in different release complexes were obtained in the same way as Fig. 1b with the rate constants k_a_ = 17 µM^-1^s^-1^, [RF2_free_] = 3 µM, k_conf_ = 11 s^-1^ and k_hydr_ = 2.7 s^-1^ (Extended Data Fig. 3). **d, e.** The populations of release complexes containing compact comformation and extended conformation of RF1 (d) and RF2 (e) at the 24 ms, 60 ms and long incubation time points as obtained by time-resolved cryo-EM after 3D classification of the particle images.

We assembled a UAA-programmed release complex, *RC_0_*, with tripeptidyl-tRNA in the P site, and visualized its structure with cryo-EM (Methods and Extended Data Fig. 1). The *RC_0_* displays no intersubunit rotation, and the tripeptide of its P-site tRNA is seen near the end of the peptide exit tunnel (Extended Data Fig. 1). The mRNA of the DC is disordered, but the overall resolution of the *RC_0_* is high (2.5Å). Apart from a small fraction of isolated ribosomal 50S subunits, the *RC_0_* ensemble is homogeneous (Extended Data Fig. 1). We used quench-flow techniques to monitor the time evolution of the class-1 RF-dependent release of tripeptide from the peptidyl-tRNA with UAA-codon in the A site after rapid mixing of RC_0_ with RF1 or RF2 at rate-saturating concentrations (k_cat_-range)^8^ (Fig. 1a). The experiments were performed at pH-values from 6 to 8 units, corresponding to [OH^-^] variation in the 0.25 to 2.5 µM range (Extended data Fig. 2 and Extended Data Fig. 3). The results are consistent with the existence of a two-step mechanism, in which a pH-independent conformational change (rate constant k_conf_) is followed by pH-dependent ester bond hydrolysis (see Methods). We estimate k_conf_ as 18 s^-1^ for RF1 and 11 s^-1^ for RF2 at 25°C, which approximates the effective incubation temperature for the time-resolved cryo-EM experiments (Extended Data Fig. 2 and Extended Data Fig. 3).

From the quench-flow data, we predicted that the ensemble fraction of the compact RF1/2 form would be predominant at 24 ms, and much smaller at 60 ms (Fig. 1b and 1c). These predictions are in qualitative agreement with the time-resolved cryo-EM data, which show a somewhat faster conformational transition than in the quench-flow experiments (Fig. 1d and 1e). We first focus on the cryo-EM structures of RF1, and then highlight the few structural differences between RF1 and RF2.

At 24 ms reaction time, 25% of ribosome-bound RF1 is in the compact form in what we name the *pre-accommodation* state of the ribosome (Fig. 1d). The 70S part of the complex is similar to that of the *pre-termination* complex preceding RF-binding, but there is an additional A-site density belonging to RF1 (Fig. 2a). In *pre-accommodation* state of the ribosome, domain III of RF1 is 60-70 Å away from the PTC, in a similar relative orientation as in the crystal forms of the free factors^1,14^ (Extended Data Fig. 4) and significantly differing from that in the post-accommodated state of the terminating ribosome^3^. The loop that contains the GGQ motif of RF1 is positioned at the side of the β-sheet of domain II (near aa 165–168), facing the anticodon-stem loop and the D stem of the P-site tRNA (Fig. 2a and 2c).

**Figure 2.**
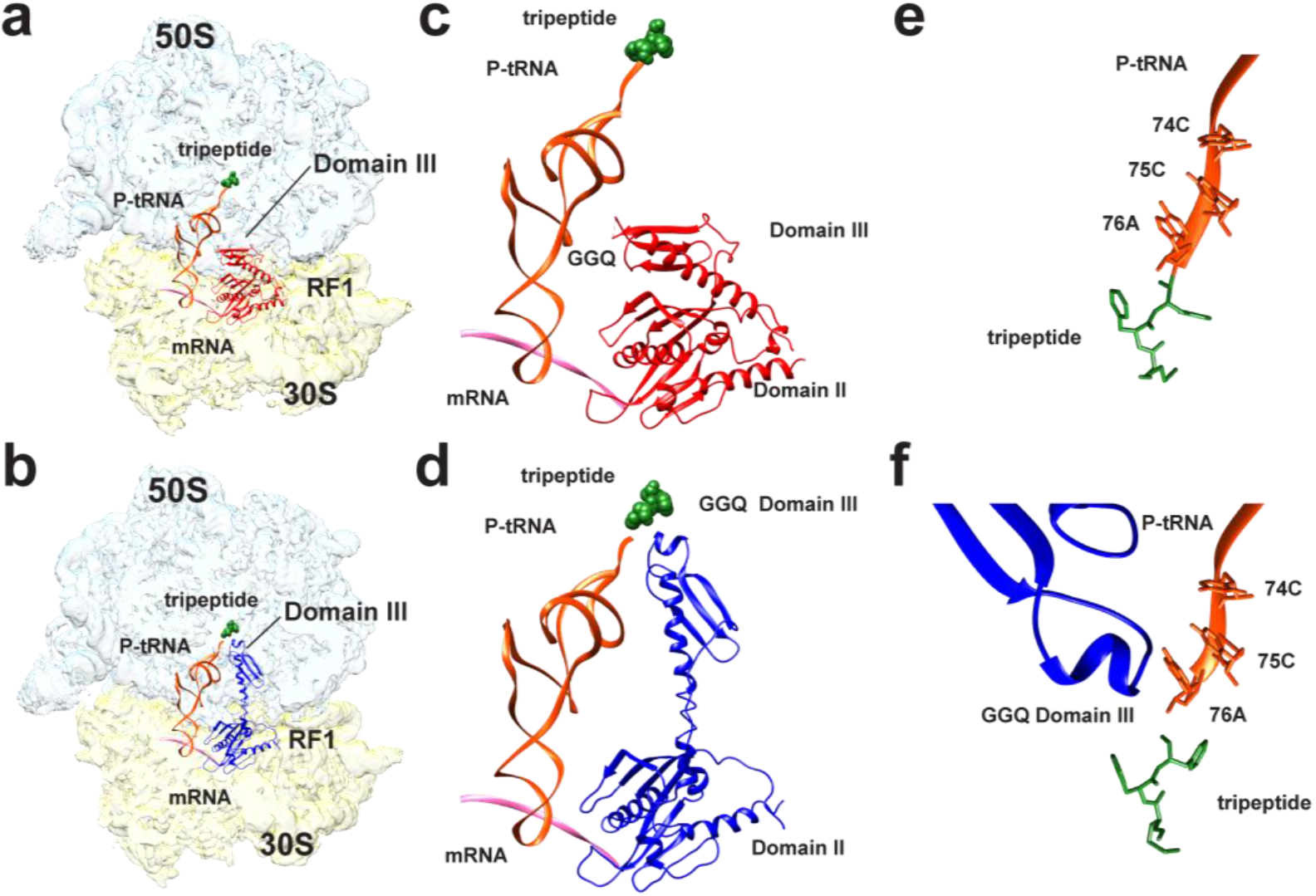
Time-resolved cryo-EM structures of *E.coli* 70S ribosome bound with release factor 1. **a.** Pre-accommodated ribosome complex bound with RF1 in a compact conformation. **b.** Accommodated ribosome complex bound with RF1 in a extended conformation. Light blue: 50S large subunit; light gold: 30S small subunit; green: tripeptide; orange: P-tRNA; pink: mRNA; red: compact RF1; and dark blue: extended RF1. **c and d.** Positions of domain III of ribosome-bound RF1 in pre-accommodated ribosome complex (c) and accommodated RF1-ribosome complex (d) relative to mRNA (pink), P-tRNA (orange) and tripeptide (green). **e and f.** Close-up views of the upper peptide exit tunnel, showing tripeptide (green) in pre-accommodated ribosome complex (e) and accommodated ribosome complex (f).

At 60 ms reaction time the RF1-bound ribosome ensemble is dominated by the extended form of RF1 (Fig. 2b and 2d). We term the ribosome complex with extended RF1 the *accommodated RF1-ribosome complex*. It contains density for the tripeptide in the exit tunnel, separated by a density gap from the CCA end of the P-site tRNA, indicating that at 60 ms the peptide has been severed from the P-site tRNA but not released from the ribosome (Fig. 2e and 2f). At a much later time-point, the tripeptide density is no longer present in the exit tunnel of the accommodated RF-ribosome complex. Precise estimation of the time evolution of tripeptide dissociation from the ribosome would require additional time points. Of particular functional relevance would be estimates of the time of dissociation of longer peptide chains from the exit tunnel after ester bond hydrolysis.

The most striking difference between the compact and extended conformation of ribosome-bound RF1 is the position of the GGQ of domain III. As RF1 switches its conformation from the compact to the extended form, the repositioning of domain III places the catalytic GGQ motif within the PTC, and adjacent to the CCA end of the P-site tRNA (Fig. 2c and 2d). The extended form of RF1 has a similar conformation as found in the previous studies^16,19,21,22,34,35^.

Similar to the case of sense-codon recognition by tRNA, three universally conserved DC residues, A1492, A1493 and G530 of the ribosome’s 16S rRNA undergo key structural rearrangements during translation termination. In the RF-lacking termination complex, A1492 of helix 44 in 16S rRNA stacks with A1913 of H69. A1493 is flipped out and stabilizes the first two bases in the A-site stop codon. G530 stacks with the third base A in the stop codon. In the presence of RF, whether compact or extended, A1492 is flipped out towards G530 and interacts with the first two stop-codon bases. A1493 stacks with A1913, which is in close contact with A1492 in the RF-lacking termination complex. G530 stacks with the third stop-codon base (Fig. 3a and 3b).

**Figure 3.**
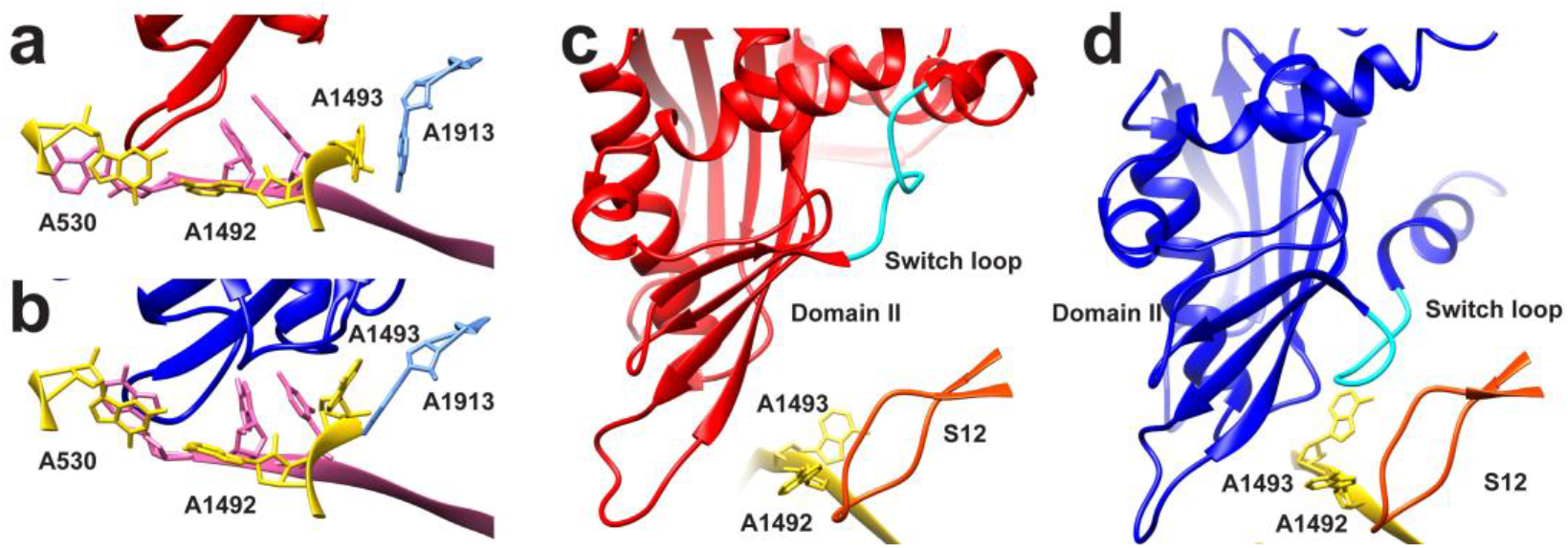
Interaction of RF1 with the ribosomal decoding center. **a and b.** Structures of the ribosomal decoding center in pre-accommodated ribosome complex (a) and accommodated ribosome complex (b). Red: compact RF1; dark blue: extended RF1. **c and d**. Conformations of switch loop in preaccommodated ribosome complex (c) and accommodated ribosome complex (d). Gold: A1492 and A1492; and orange: S12.

The switch loop, which was previously proposed to be involved in inducing a conformational change of RF1^19,35^, shows no interaction with protein S12 or 16S rRNA in the compact form of RF1 (Fig. 3c) whereas in the extended form of RF1, the rearranged conformation of the switch loop is stabilized by interactions within a pocket formed by protein S12, the loop of 16S rRNA, the β-sheet of domain II, and the flipped-out nucleotides A1492 and A1913 (Fig. 3d). Shortening the switch loop (302-304) resulted in a substantially slower, rate-limiting step in peptide release^36^, which indicates that the switch loop plays a role in triggering the conformational change of RF1.

Similar experiments were carried out for RF2 at 24 ms, 60 ms and 5 h reaction times. RF2 undergoes a conformational change from compact to expanded form similar to that of RF1 (Fig. 1e). As in the case of RF1, the switch loop of RF2 makes no contact with protein S12 or 16S rRNA. In the extended form of RF2, A1492 is flipped out from helix 44 (h44) of 16S rRNA and stacks on the conserved Trp319 of the switch loop, stabilizing the extended conformation of RF2 on the ribosome.

The ribosome complexes with RF1/2 bound in compact conformation seen here are distinct from those reported for the ribosome rescue complex, in which ArfA is bound in the A site lacking a stop codon^5,6^. In our structures, the conformation of the conserved 16S rRNA residues in the DC (A1492, A1493 and G530) is similar to the classical termination configuration^19^. In contrast, in the presence of ArfA, these residues adopt conformations known from sense-codon recognition^5,6^. It suggests that the compact RFs bind to the A site regardless of the conformation of the DC. The conformational change of RFs is likely due to the changes in the switch loop triggered by its interaction with protein S12 and 16S rRNA. This interaction is disrupted by the mutation A18T of ArfA, hence leaving RFs in the compact conformation^6^.

Our ribosome complexes with RF1/RF2 are also distinct from a recent ribosome complex with compact RF1, reported by Korostelev and collaborators ^7^. Shortening of the switch loop, combined with the addition of the antibiotic Bl-S which prevents the GGQ motif from reaching the PTC, stabilizes ribosome-bound RF1 in a compact conformation^7^, distinct from the transient, compact RF1-structure observed here. In our structure, the SCR between the β4–β5 strands on domain II are bound loosely to the A site (Fig. 3a). In the Bl-S-halted compact RF1 structure^7^, in contrast, the stop codon-recognition motif of RF1 has moved further into the A site by 5 Å, to a position almost identical to that of the fully accommodated, extended structure of RF1. The functional role of their structure is not immediately obvious, but if it can be interpreted as an authentic transition state analogue, the roles of our respective complexes would be complementary. We previously found that high accuracy of stop signal recognition depends on smaller dissociation constant (K_m_-effect) and larger catalytic rate constant (k_cat_-effect) for class-1 RF reading of cognate stop codons compared to near-cognate sense codons^37^. The K_m_-effect contributes by factors from 100 to 3000 and the k_cat_-effect by factors from 2 to 3000 to the overall termination accuracy values in the 10^3^ to 10^6^ range ^37^. Accordingly, the present structure may represent binding of RFs in a transient state whereg rapid and codon-selective dissociation rates are responsible for the accuracy factor due to the K_m_-effect. Furthermore, Korostelev’s structure ^7^, with its comparatively deep interaction between the cognate stop codon and SCR center, could mimic the authentic transition state on the path from compact to the extended form of the RF. Accordingly, Korostelev’s structure may illustrate additional selectivity due to the k_cat_-effect. To test these hypotheses, molecular computations^27^ based on our respective RF structures could be used to compare their stop codon selectivities with those of RFs in the post-termination state of the ribosome^27^.

In conclusion, during translation termination, the release of the nascent peptide must be strictly coordinated with the recognition of a stop codon at the A site. Our cryo-EM analysis shows that in the presence of a class-1 RF the bacterial ribosome adopts several states. After rapid addition of RF1 or RF2 to a ribosomal termination complex with tripeptidyl-tRNA attached at the P site, we first observe the pre-accommodated RF-ribosome complex (compact form of RF) at 20 ms with the peptide still attached to the P-site tRNA. This, we suggest, is the first step in the termination reaction. Second, at 60 ms, we observe the accommodated RF-ribosome complex with the extended form of RF and the tripeptide now in the exit tunnel and no longer attached to the P-site tRNA. Third, at a much later time point, we observe the post-accommodated RF-ribosome complex, with the extended form of RF without tripeptide in the exit tunnel (Extended Data Fig. 5). These pieces of evidence from our time-resolved experiments clearly reflect the sequence of events in termination of bacterial protein synthesis. A structure-based model for the stepwise interaction between ribosome and RF and the release of the nascent peptide from the termination complex during the translation termination process is presented in Fig. 4. It shows how the ribosome traverses (1) the pre-termination state with the stop codon at the A site, (2) the initial binding state (RF compact; “pre-accommodated RF-ribosome complex”), (3) the open catalytic state (RF open/extended; “accommodated RF-ribosome complex”) and (4) the state after peptide release. Finally, we suggest that the selective advantage of the compact RF-form is that it allows for rapid factor binding into and dissociation from an accuracy maximizing pre-accommodation state.

**Figure 4.**
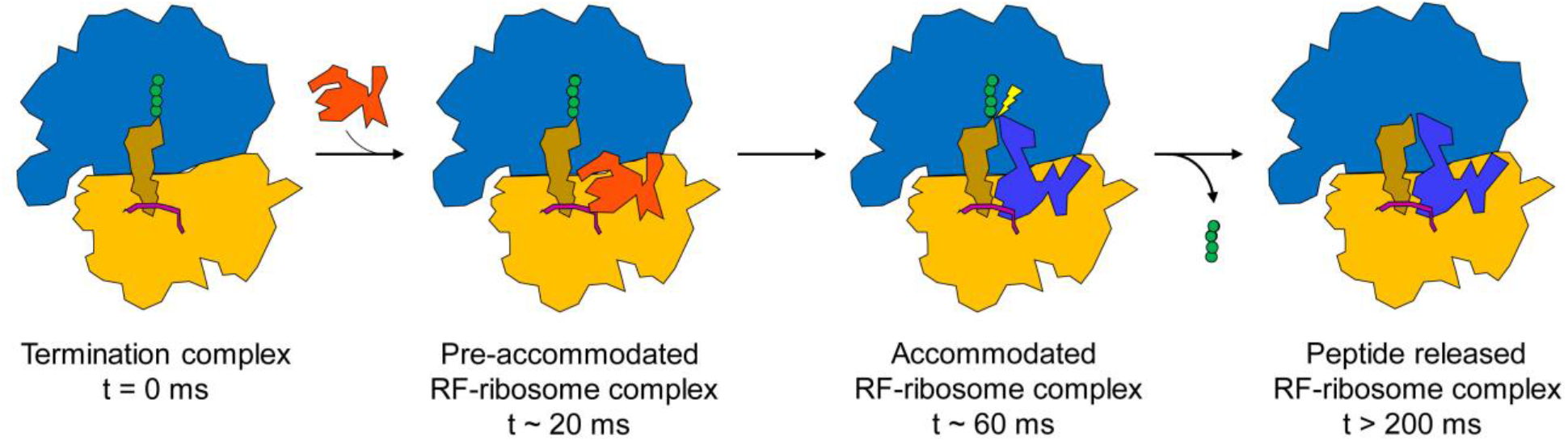
Structure-based model for the stepwise interaction between ribosome and RF and the release of the nascent peptide from the termination complex during the translation termination process. The sequence of states is (1) the termination complex with the stop codon at the A site, (2) the initial binding state (RF compact; “pre-accommodated RF-ribosome complex”), (3) the open catalytic state (RF open/extended; “accommodated RF-ribosome complex”) and (4) the state after peptide release. Blue: 50S large subunit; orange: 30S small subunit; green: tripeptide; brown: P-tRNA; pink: mRNA; red: compact RF; and blue-purple: extended RF.

## Methods

### Preparation of components for cell-free protein synthesis and fast kinetics experiments

Buffers and all *Escherichia coli (E. coli)* components for cell-free protein synthesis were prepared as described^8^. Ribosomal release complexes (RC) contained tritium (^3^H) labelled fMet-Phe-Phe-tRNA^Phe^ in the P site and had UAA stop-codon programmed A site. The mRNA sequence used to synthesize the peptide was GGGAAUUCGGGCCCUUGUUAACAAU<u>UAAGGAGG</u>UAUUAA**AUGUUCUUCUAA**UGCAGAAAAAAAAAAAAAA AAAAAAA (ORF in bold, SD underlined). Class-1 release factors (RFs), overexpressed in *E. coli*, had mainly unmethylated glutamine (Q) in the GGQ motif and the RF2 variant contained Ala in position 246. Rate constants for conformational changes of RFs in response to cognate A-site stop codon (k_conf_) and for ester bond hydrolysis (k_hydr_) at different OH^-^ concentrations were estimated as described^8^. In short, purified release complexes (0.02 µM final concentration) were reacted at 25°C with saturating amounts of RFs (0.8 µM final) in a quench-flow instrument at, and the reaction stopped at different time points by quenching with 17% (final concentration) formic acid. Precipitated [^3^H]fMet-Phe-Phe-tRNA^Phe^ was separated from the soluble [^3^H]fMet-Phe-Phe peptide by centrifugation. The amounts of tRNA-bound and free peptides were quantified by scintillation counting of the ^3^H radiation. Reaction buffer was polymix-HEPES with free Mg^2+^ concentration adjusted from 5 mM to 2.5 mM by addition of 2.5 mM Mg^2+^-chelating UTP. The rate constants for RF association to the A site at 25°C, *k_a25_*, were estimated from their previously published values at 37°C, *k_a37_ = 60 µM^-1^s^-1^* for RF1 and 23 *µM^-1^s^-1^* for RF2^37^ through ka25 =(T_25_ /ŋ_25_)·(ŋ_37_ /T_37_), where T is the absolute temperature and ŋ the water viscosity. Kinetics simulations were carried out with the termination reaction steps modelled as consecutive first-order reactions^38^.

### Preparation of EM grids and mixing-spraying time-resolved cryo-EM

Quantifoil R1.2/1.3 grids with a 300 mesh size were subjected to glow discharge in H_2_ and O_2_ for 25 s using a Solarus 950 plasma cleaning system (Gatan, Pleasanton, CA) set to a power of 10 W. Release complexes and RFs were prepared in the same way as for quench-flow experiments, except the release complexes were unlabeled. For each time point (24 ms and 60ms), 4 µM of release complexes in polymix-HEPES with 2.5 mM UTP and 6 µM of class-1 release factors in the same buffer were injected into the corresponding microfluidic chip at a rate of 3 µl/s such that they could be mixed and sprayed onto a glow-discharged grid as previously described ^12^. The final concentration of the release complexes and the class-1 release factors after rapid mixing in our microfluidic chip was 2 µM and 3 µM, respectively. As the mixture was sprayed onto the grid, the grid was plunge-frozen in liquid ethane-propane mixture and stored in liquid nitrogen until it was ready to be imaged.

### Preparation of EM grids and blotting-plunging cryo-EM

Grids of *RC_0_* and long-incubation complex were prepared with the following protocol. 3uL sample was applied in the holey grids (gold grids R0.6/1 300 mesh, which was plasma cleaned using the Solarus 950 advanced plasma cleaning system (Gatan, Pleasanton, CA) for 25 s at 10 W using hydrogen and oxygen plasma). Vitrification of samples was performed in a Vitrobot Mark IV (FEI company) at 4 °C and 100% relative humidity by blotting the grids once for 6 s with a blot force 3 before plunging them into the liquid alkane.

### Cryo-EM data collection

Time-resolved cryo-EM grids were imaged either with a 300 kV Tecnai Polara F30 TEM or a Titan Krios TEM. The images were recorded at a defocus range of −1 to −3 µm on a K2 direct detector camera (Gatan, Pleasanton, CA) operating in counting mode with pixel size at 1.66 Å or 1.05 Å. A total of 40 frames were collected with an electron dose of 8 e^−^/pixel/s for each image. Blotting-plunging cryo-EM grids were imaged with a 300 kV Tecnai Polara F30 TEM. The images were recorded at a defocus range of 1-3 µm on a K2 direct detector camera (Gatan, Pleasanton, CA) operating in counting mode with pixel size at 1.24 Å. A total of 40 frames were collected with an electron dose of 8 e^−^/pixel/s for each image.

### Cryo-EM data processing

The beam-induced motion of the sample and the instability of the stage due to thermal drift was corrected using the MotionCor2 software program^39^. The contrast transfer function (CTF) of each micrograph was estimated using the CTFFIND4 software program^40^. Imaged particles were picked using the Autopicker algorithm included in the RELION 2.0 software program ^41^. For each time point (Extended Data Fig. 6 and Extended Data Fig. 7), 2D classification of the recorded images were used to separate 70S ribosome-like particles from ice-like and/or debris-like particles picked by the Autopicker algorithm and to classify the particles that were picked for further analysis into 70S ribosome-like particle classes. These particle classes were then combined into a single dataset of 70S ribosome-like particles and subjected to a round of 3D classification for the purpose of eliminating those obvious contaminants from the rest of the dataset. This classification was set for 10 classes with the following sampling parameters: Angular sampling interval of 15°, offset search range of 5 pixels and offset search step of 1 pixel. The sampling parameters were progressively narrowed in the course of the 50 classification iterations, down to 3.7° for the angular sampling interval. At the end of the first classification round, two classes were found inconsistent with the known structure of the 70S ribosomes and were thus rejected. The rest of the particles were regrouped together as one class. All particles from this class were re-extracted using unbinned images. A consensus refinement was calculated using these particles. The A site of the 70S ribosome displays fractioned density indicating heterogeneity, then therefore the signal subtraction approach was applied. The A-site density was segmented out of the ribosome using Segger in Chimera^42^. The mass of density identified as release factor was used for creating a mask in RELION with 3 pixels extension and 6 pixels soft edge using relion_mask_create. This mask was used for subtracting the release factor-like signal from the experimental particles. The new particles images were used directly as input in the masked classification run with the number of particles set for ten classes, and with the mask around the release factor-binding region. This run of focused classification resulted in two separate classes, one with compact and one with extended conformation of the release factors. The corresponding raw particles were finally used to calculated consensus refinements. The local resolution of the final maps was computed using ResMap^43^.

For the *RC_0_* complex dataset, the software MotionCor2^39^ was used for motion correction and dose weighting. Gctf^44^ was used for estimation of the contrast transfer function parameters of each micrograph. RELION^41^ was used for all other image processing steps. Particles picking was done automatically in RELION. Boxed out particles were extracted from dose-weighted micrographs with eight times binning. 2D classifications were initially performed on bin8 particle stacks to remove false positive particles from the particle picking step. 3D classification were performed on bin4 particle stacks. Classes from bin4 and bin2 3D classification showing high-resolution features were saved for further processing steps. Un-binned particles from this class were re-extracted and subjected to auto-refinement. The final density map was sharpened by applying a negative B-factor estimated by automated procedures. Local resolution variations were estimated using ResMap^43^ and visualized with UCSF Chimera ^42^.

### Model building and refinement

Models of the *E. coli* 70S ribosome (5MDV, 5MDW and 5DFE) were docked into the maps using UCSF Chimera. The pixel size was calibrated by creating the density map from the atomic model and changing the pixel size of the map to maximize the cross-correlation value. For the compact RF1 model, a homology model was generated with the crystal structure of the RF1 (PDB ID: 1ZBT) as a template using the SWISS-MODEL online server^45^. This homology model was rigid-body-fitted into the map using UCSF Chimera, followed by manual adjustment in Coot^46^. Due to the lack of density, domain I of RF1 was not modelled.

### Figure preparation

All figures showing electron densities and atomic models were generated using UCSF Chimera^42^.

### Data availability

The data that support the findings of this study are available from the corresponding author upon request. The atomic coordinates and the associated maps have been deposited in the PDB and EMDB with the accession codes.

## Acknowledgements

This work was supported by HHMI and grants NIH R01 GM55440 and R01 GM29169 (to J.F.), the Swedish Research Council, and the Knut and Alice Wallenberg Foundation (to M.E.), and the Sederholms travel stipend (Uppsala University) (to G.I.).

## Author contributions

Z.F., G.I., S.K., B.S., M.S., B.C., R.A.G., M.E. and J.F. conceived and designed experiments. G.I. and M.E. carried out biochemical experiments. Z.F., S.K., G.I., and B.S. performed time-resolved cryo-EM experiments. Z.F., S.K., G.I., B.S., and M.S. performed image processing and atomic modelling. Z.F., S.K., G.I., and B.S. analysed the data. Z.F., G.I., S.K., M.E. and J.F. wrote the manuscript.

---

**Extended Data Fig. 1.**
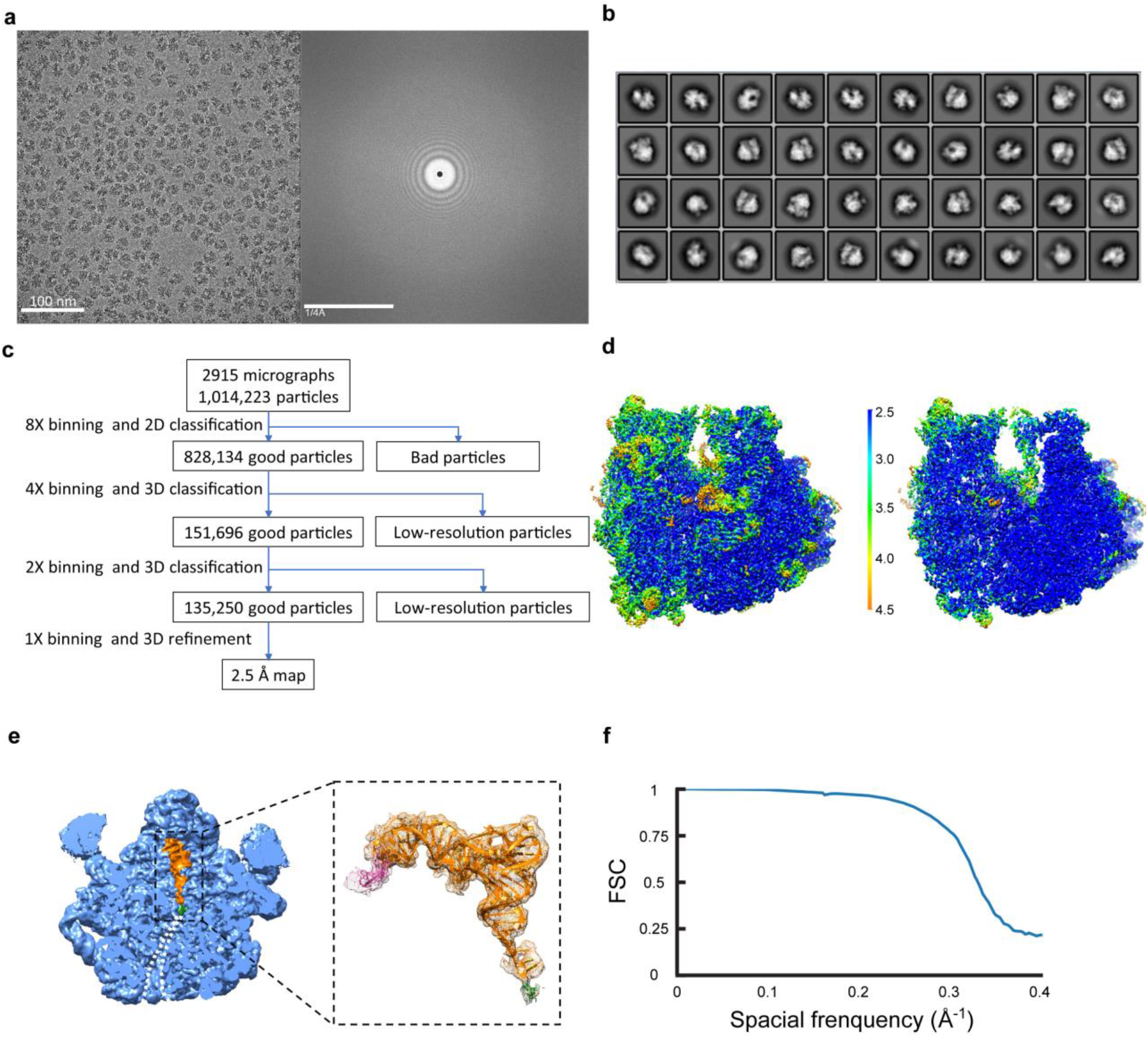
Cryo-EM data processing of release complex, *RC_0_*. **a.** A representative micrograph collected with a Polara transmission electron microscope with corresponding power spectrum. **b.** Representative 2D class averages from reference-free 2D classification. **c.** Particle classification and structural refinement procedures used. **d.** Local resolution estimation of the cryo-EM density map. The density map of release complex is displayed in surface representation, and colored according to the local resolution (see color bar). **e.** The ribosomal exit tunnel (white dash line) in 50S ribosomal subunit (blue) is occupied by programed tripeptide (green) attached to P-site tRNA (orange). The interaction between mRNA (pink) and tRNA is shown in zoom-in view on the right. **f.** FSC curve for cryo-EM reconstruction.

**Extended Data Fig. 2.**
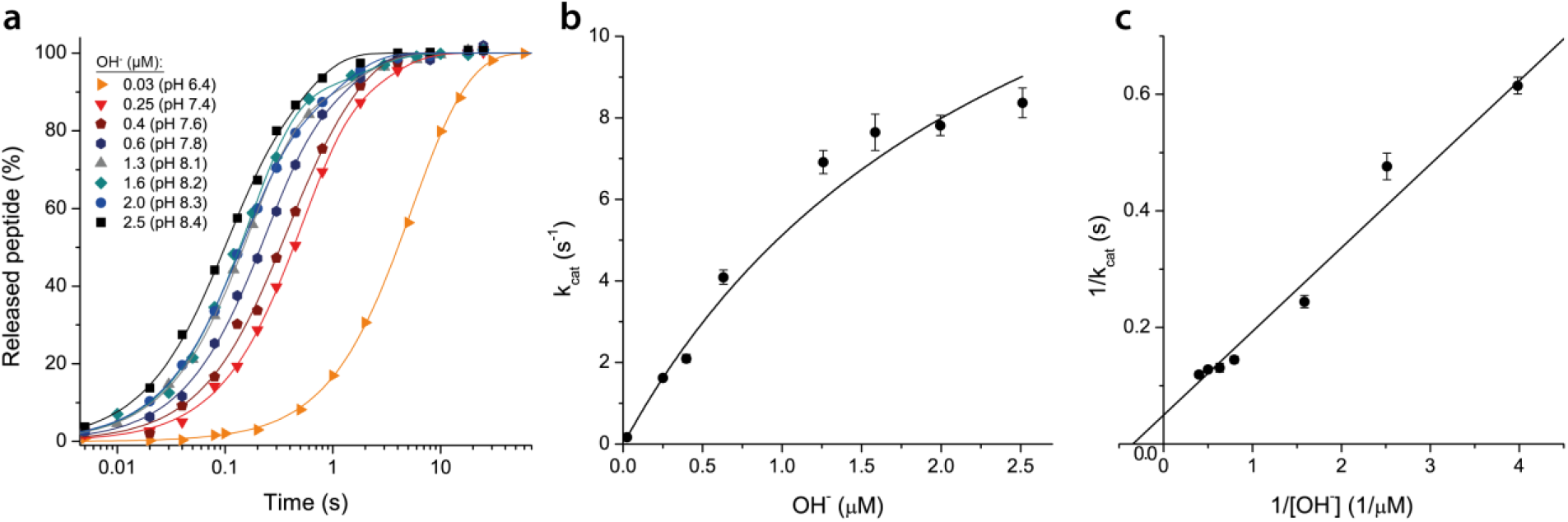
The maximal rate (k_cat_) of peptide release by RF1 has a pH-independent and a pH-dependent step. **a.** Extent of RF1-dependent fMet-Phe-Phe peptide release from P-site bound fMet-Phe-Phe-tRNA (y axis) versus time (x axis) at different hydroxyl ion (OH^-^) concentrations. The experiments were performed at 25°C with 0.02 µM release complexes and saturating (0.8 µM) RF1 concentration (k_cat_-range) (Methods). **b.** Maximal rate (k_cat_) of peptide release (y-axis) increased to a plateau value (kconf = 18.1 ± 3.3 s^-1^) with increasing [OH^-^] (x-axis), which led to the hypothesis that the maximal rate of peptide release is limited by a pH-independent change of RF conformation at high pH^8^. Error bars represent s.d. from three replicates. The Michaelis-Menten model, used to obtain k_conf_, was compared to a linear model (k_cat_ linearly increasing with OH^-^ concentration) using the F-test. The linear model could be rejected (F_5,6_ = 35, p < .001). **c.** Lineweaver-Burk plot of 1/k_cat_ versus 1/[OH^-^] from the data in Extended Data Fig. 2b. The value at 1/0.025 µM OH^-^ (x = 40; y = 6) is not shown for better visibility of the remaining data points. The plateau value k_conf_ was obtained from the plot including all values as 1/y-intercept and was 16.7 ± 3.6 s^-1^.

**Extended Data Fig. 3.**
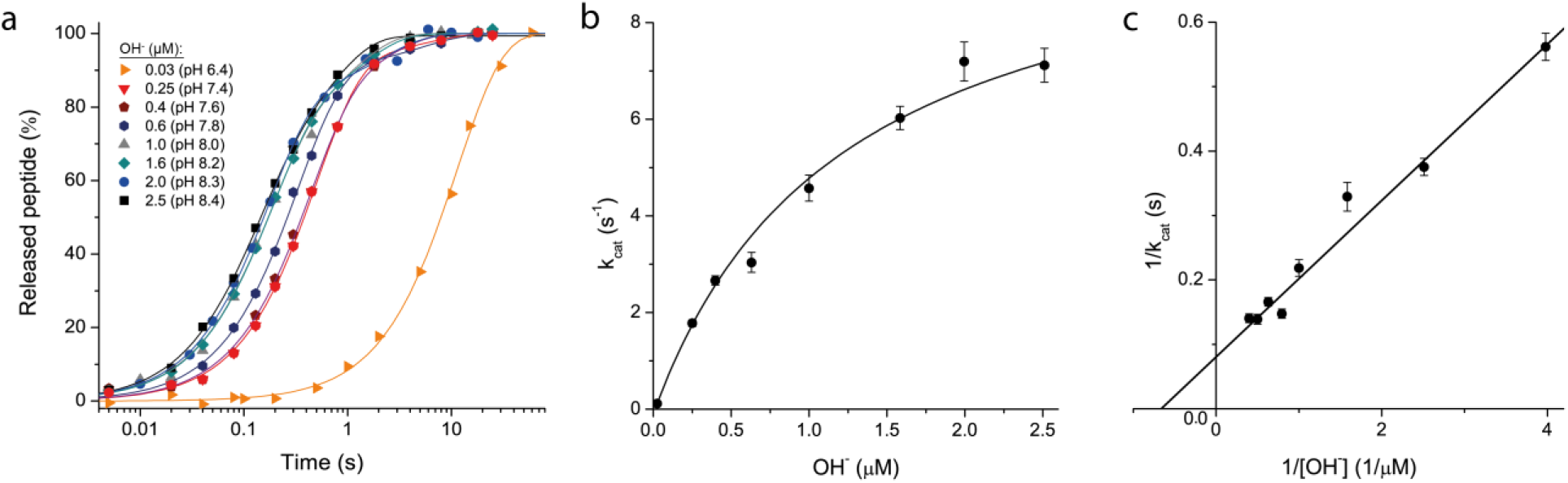
The maximal rate (k_cat_) of peptide release by RF2 has a pH-independent and a pH-dependent step. **a.** Extent of RF2-dependent fMet-Phe-Phe peptide release from P-site bound fMet-Phe-Phe-tRNA (y axis) versus time (x axis) at different hydroxyl ion (OH^-^) concentrations. The experiments were performed at 25°C with 0.02 µM release complexes and saturating (0.8 µM) RF2 concentration (Methods). **b.** Maximal rate (k_cat_) of peptide release (y-axis) by RF2 increased to a plateau value (kconf = 10.8 ± 0.9 s^-1^) with increasing [OH^-^] (x-axis). Error bars represent s.d. from three replicates. The Michaelis-Menten model, used to obtain k_conf_, was compared to a linear model (k_cat_ linearly increasing with OH^-^ concentration) using the F-test. The linear model could be rejected (F_5,6_ = 141, p < .001). **c.** Lineweaver-Burk plot of 1/k_cat_ versus 1/[OH^-^] from the data in Extended Data Fig. 3b. The value at 1/0.025 µM OH^-^ (x = 40; y = 8.5) is not shown for better visibility of the remaining data points. The plateau value k_conf_ was obtained from the plot including all values as 1/y-intercept and was 12.5 ± 1.2 s^-1^.

**Extended Data Fig. 4.**
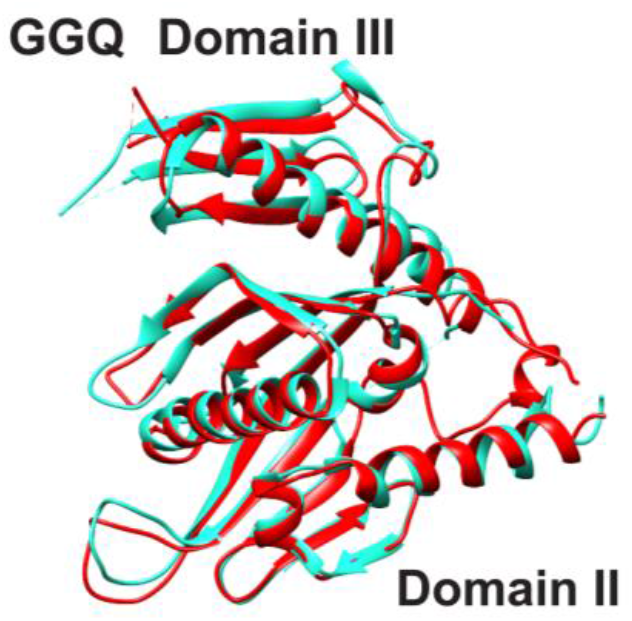
Comparison of compact RF1 from pre-accommodated ribosome complex (red) and free RF1 in the crystal form (PDB 1RQ0).

**Extended Data Fig. 5.**
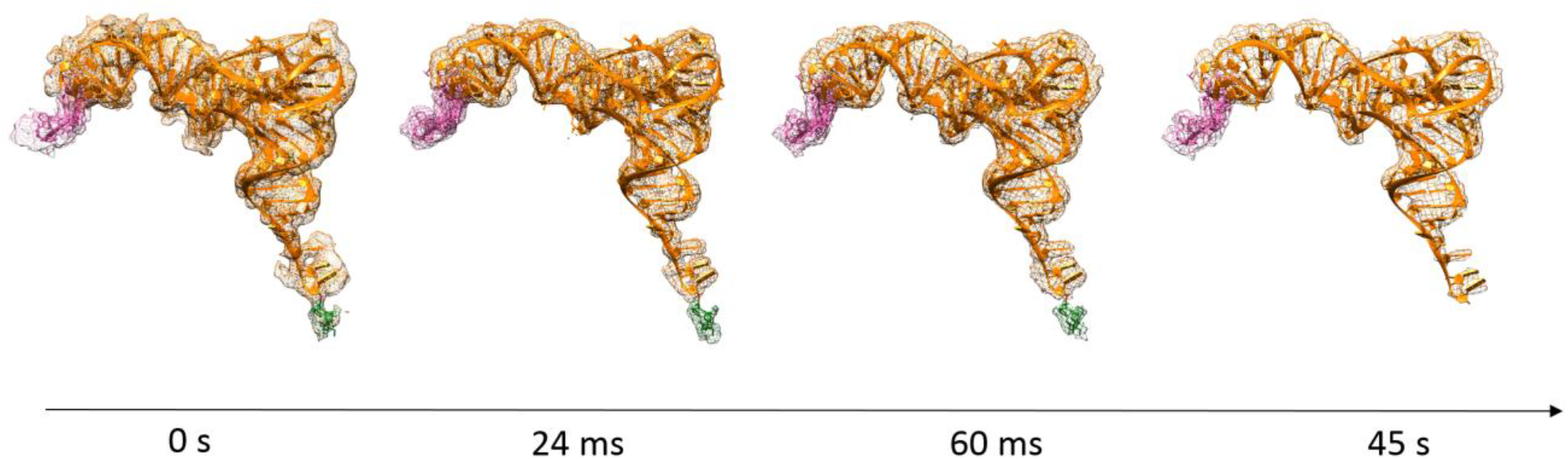
Tripeptide hydrolysis and release. mRNA, P-tRNA and tripeptide density with refined models of RC_0_ from 0 s, compact RF1 bound pre-accommodated ribosome from 24 ms, extended RF1 bound accommodated ribosome from 60 ms, and extended RF1 bound accommodated ribosome from 45 s, respectively (left to right). Pink: mRNA; orange: tRNA; and green: tripeptide.

**Extended Data Fig. 6.**
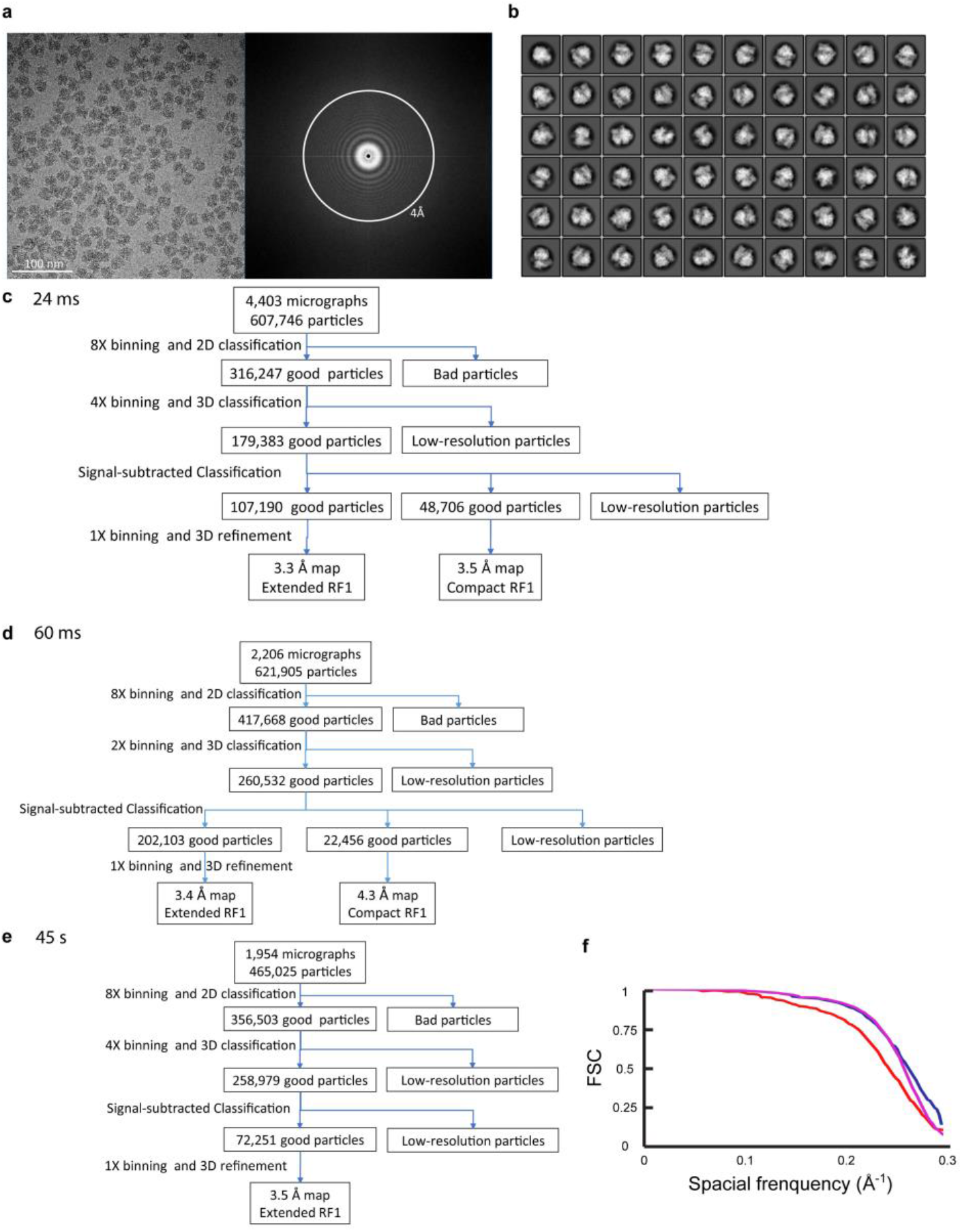
Cryo-EM data processing of RF1-bound ribosome ensembles at 24 ms, 60 ms, and 45s. **a.** A representative micrograph with corresponding power spectrum. **b.** Representative 2D class averages from reference-free 2D classification. **c,d, and e.** Particle classification and structural refinement procedures used for 24 ms (c), 60 ms (d), and 45 s (e) data sets. **f.** FSC curves for cryo-EM reconstructions of compact RF1 bound pre-accommodated ribosome from 24 ms (red), extended RF1 bound accommodated ribosome from 60 ms (blue), and extended RF1 bound accommodated ribosome from 45 s (purple), respectively.

**Extended Data Fig. 7.**
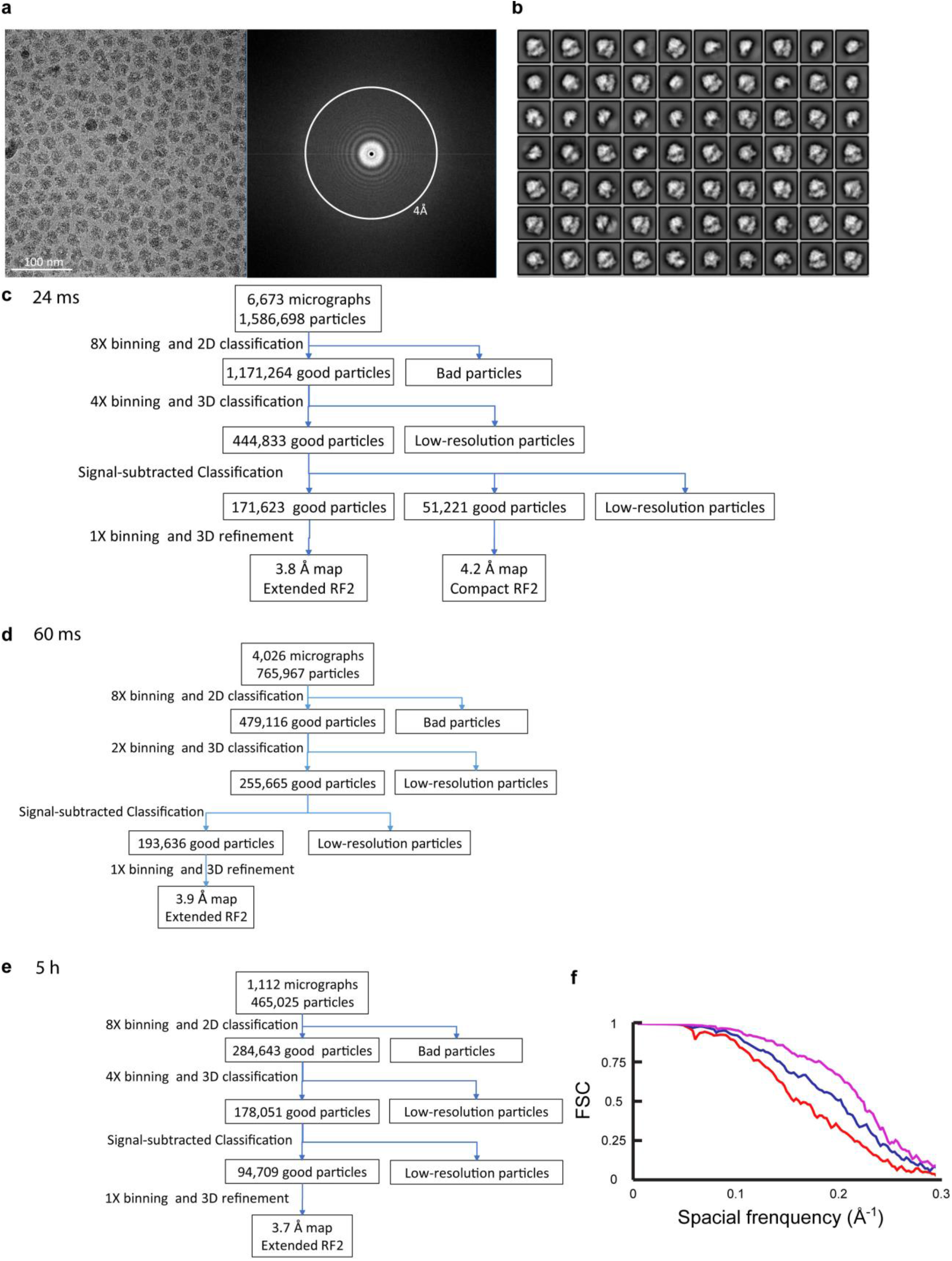
Cryo-EM data processing of RF2-bound ribosome ensembles at 24 ms, 60 ms, and 5h. **a.** A representative micrograph with corresponding power spectrum. **b.** Representative 2D class averages from reference-free 2D classification. **c,d, and e.** Particle classification and structural refinement procedures used for 24 ms (c), 60 ms (d), and 5 h (e) data sets. **f.** FSC curves for cryo-EM reconstructions of compact RF2 bound pre-accommodated ribosome from 24 ms (red), extended RF2 bound accommodated ribosome from 60 ms (blue), and extended RF2 bound accommodated ribosome from 5 h (purple), respectively.

**Extended Data Table 1.** Cryo-EM data collection and model statistics of ribosome complexes.

|  | release complex | pre-accommodated | accommodated | accommodated | pre-accommodated | accommodated | accommodated |
| --- | --- | --- | --- | --- | --- | --- | --- |
| **Data Collection** |  |  |  |  |  |  |  |
| Electron microscope | Polara | Krios | Polara | Polara | Polara | Polara | Polara |
| Pixel size (Å) | 1.24 | 1.04 | 1.66 | 1.66 | 1.66 | 1.66 | 1.66 |
| Particle number | 135250 | 48706 | 202103 | 72251 | 51221 | 193636 | 94709 |
| Time point | 0s | 24 ms | 60 ms | 45 s | 24 ms | 60 ms | 5h |
| Release factor | unbound | RF1 | RF1 | RF1 | RF2 | RF2 | RF2 |
| Map resolution (Å) | 2.5 | 3.5 | 3.4 | 3.5 | 4.2 | 3.9 | 3.7 |
| **Refinement** |  |  |  |  |  |  |  |
| Model-to-map | 0.9106 | 0.8697 | 0.8485 | 0.8695 | 0.8762 | 0.8719 | 0.8471 |
| **R.M.S Deviation** |  |  |  |  |  |  |  |
| Bond lengths (Å) | 0.009 | 0.009 | 0.008 | 0.008 | 0.009 | 0.006 | 0.008 |
| Bond angles | 0.801 | 0.881 | 0.843 | 0.877 | 0.999 | 0.822 | 0.937 |
| **Ramachandran plot** |  |  |  |  |  |  |  |
| Outliers | 0.04% | 0.03% | 0.05% | 0.03% | 0.03% | 0.03% | 0.03% |
| Allowed | 3.59% | 6.09% | 5.27% | 5.56% | 6.86% | 5.08% | 6.19% |
| Favored | 96.38% | 93.88% | 94.68% | 94.41% | 93.11% | 94.88% | 93.78% |
| Rotamer outliers | 0.43% | 0.19% | 0.37% | 0.41% | 0.18% | 0.18% | 0.49% |
| Cbeta deviations | 0% | 0% | 0% | 0% | 0% | 0% | 0% |

